# Epileptiform activity influences theta-burst induced LTP in the adult hippocampus: a role for lipid raft dynamics in early metaplasticity

**DOI:** 10.1101/2022.12.06.519267

**Authors:** JD Carvalho-Rosa, NC Rodrigues, A Silva-Cruz, SH Vaz, D Cunha-Reis

## Abstract

Non-epileptic seizures are identified as a common epileptogenic trigger. Early metaplasticity following seizures may contribute to epileptogenesis by abnormally altering synaptic strength and homeostatic plasticity. We now studied how *in vitro* epileptiform activity triggers early changes in CA1 long-term potentiation (LTP) induced by theta-burst stimulation (TBS) in rat hippocampal slices and the involvement of lipid rafts in these early metaplasticity events. Two forms of epileptiform activity (EA) were induced: 1) interictal-like EA triggered by withdrawal of Mg^2+^ and K^+^ elevation to 6mM in the superfusion medium or 2) ictal-like EA induced by bicuculine (10μM) delivery. LTP induced 30 min post EA was impaired, an effect more pronounced after ictal-like EA. LTP recovered to control levels 60 min post interictal-like EA but was still impaired 60 min after ictal-like EA. The synaptic molecular events underlying this altered LTP were investigated 30 min post EA. Synaptosomes isolated from parallel slices showed enhanced AMPA GluA1 Ser831 phosphorylation but decreased Ser 845 phosphorylation and a marked decrease in GluA1/GluA2 ratio. A marked decrease in flotillin-1 and caveolin-1 levels concomitantly with a moderate increase in PSD-95 and marked increase in gephyrin levels.

Altogether, EA differentially influences hippocampal CA1 LTP thorough regulation of GluA1/GluA2 levels and AMPA GluA1 phosphorylation suggesting altered LTP post-seizures may be a relevant target of antiepileptogenic therapies. In addition, this metaplasticity is also associated with marked alterations in classic and synaptic lipid raft markers, suggesting these may also constitute promising targets in epileptogenesis prevention.

## Introduction

Epilepsy, a multifaceted disease characterized by the development of recurrent, unprovoked seizures, is a neurological disorder that affects over 65 million people worldwide. Its high-incidence and high-prevalence (6.4 cases per 1,000 persons) is nevertheless minored by its high mortality owing to accidents, *status epilepticus* or sudden unexpected death in epilepsy, often occurring during seizures (Devinsky et al., 2018). Being the most frequent, chronic, severe neurological disease it is a huge burden for public health systems (Moshé et al., 2015; Devinsky et al., 2018; Vezzani et al., 2019; Cunha-Reis et al., 2021) and since over one third of epilepsy cases are refractory to treatment with multiple antiepileptic drugs (AEDs), it is imperative to identify new therapeutic targets.

Epileptogenesis, the process by which healthy brain networks develop enhanced susceptibility to generate spontaneous recurrent seizures leading to the development of epilepsy (Pitkänen et al., 2015; Vezzani et al., 2019), is currently perceived as a continuous and progressive process extending far beyond the latent period. Yet, the early pathophysiological mechanisms involved in epileptogenesis are still for the most part unknown and tackling epileptogenesis is an unmet clinical need (Pitkänen et al., 2015; Cunha-Reis et al., 2021). One key element is that different triggers likely lead to different seizure-onset, which will ultimately determine its first line of treatment (Sloviter, 2017). As such, establishing therapeutic targets in this early phase of epileptogenesis requires a better knowledge of these mechanisms.

Mesial temporal lobe epilepsy (MTLE) involves seizures typically arising in the hippocampus or other mesial temporal lobe structures. Only about 11-26% of patients with MTLE-HS achieve complete seizure control under pharmacological treatment with existing AEDs. As such, patients often require neurosurgical resection of the epileptic focus as a last resource to ameliorate seizure recurrence and to prevent fast worsening of the disease (Liu et al., 2012; Kuang et al., 2014; Thom, 2014). The aetiology of MTLE epileptogenesis in still unknown, yet putative precipitating events such as trauma, complex febrile seizures, status epilepticus, inflammatory insults, or ischemia have been implicated. Such events may trigger epileptogenesis by generating aberrant synaptic plasticity/neuronal excitability, excitotoxicity, secondary non-convulsive *status epilepticus,* neuroinflammation and generation of reactive oxygen species that develop during a silent period when spontaneous seizures do not occur (Thom, 2014; Gambardella et al., 2016; Sloviter, 2017) but their exact chronology it is not often easy to establish partly due to the multitude of putative epileptogenic events.

Within the first days of disease progression, enhanced neurogenesis and axonal sprouting accompany changes in neuronal excitability and synaptic plasticity driving the formation of novel synaptic connections and aberrant neural networks (Beck and Yaari, 2008). Synaptic plasticity, the ability of synapses to undergo long-lasting, activity-dependent bidirectional changes in the strength of synaptic communication, can be mediated by altering the gain of stimulus-secretion coupling at the presynaptic component or by changes in the type, number, or properties of the neurotransmitter receptors and their coupling to the intracellular signalling machinery at the postsynaptic level (Bliss and Collingridge, 2019; Cunha-Reis and Caulino-Rocha, 2020). It has been hypothesized that such changes may occur maladaptively immediately following non-epileptic seizures, thus triggering epileptogenesis (Avanzini et al., 2014), but this has not so far been thoroughly investigated.

Lipid rafts are membrane microdomains involved in synaptic receptor clustering, synaptic signalling, synaptic vesicle recycling and neurotransmitter release (Allen et al., 2007) and may have a crucial role in both synaptic and homeostatic plasticity under such pathological conditions (Sebastião et al., 2013). Synaptic lipid raft markers include both classic raft-associated proteins like caveolin-1 and flotillin-1, characteristic of caveolae and planar lipid rafts, respectively, and structural postsynaptic receptor anchoring proteins like PSD-95, present at glutamatergic synapses, and gephyrin, distinctive of GABAergic synapses (Tulodziecka et al., 2016; Papadopoulos et al., 2017; Bai et al., 2021). The latter two are also recognized as important NMDA, AMPA and GABA_A_ anchoring proteins, respectively.

*In vitro* models of ictogenesis have provided important insights into the therapeutic potential of several candidate anti-seizure drugs (Gloveli et al., 1995; Antonio et al., 2016; Raimondo et al., 2017; Dulla et al., 2018). In this paper we used such *in vitro* models of epileptiform activity to evaluate the time-course of changes in LTP occurring within 30 min to 1h 30m following seizures. LTP was evoked *in vitro* by theta burst stimulation (TBS), a sequence of electrical stimuli that mimic CA1 pyramidal cell burst firing as occurring during the hippocampal theta rhythm (4-10Hz), an EEG pattern linked to hippocampal memory storage in rodents and involves the recruitment of GABAergic mechanisms (Vertes, 2005; Larson and Munkácsy, 2015; Artinian and Lacaille, 2018; Rodrigues et al., 2021). We characterized also the synaptic molecular alterations in AMPA receptors involved in these early alterations. In addition, its dependency on lipid raft dynamics, an important requirement for several molecular mechanisms involved in synaptic plasticity, was also investigated. A preliminary account of some of the results has been published as an abstract.

## Methods

All animal procedures and protocols were performed according to ARRIVE guidelines for experimental design, analysis, and their reporting. Animal housing and handing was performed in accordance with the Portuguese law (DL 113/2013) and European Community guidelines (86/609/EEC and 63/2010/CE). The experiments were performed on hippocampal slices (400 μm thick), cut perpendicularly to the long axis of the hippocampus, obtained from young-adult (6-7 weeks old) male outbred Wistar rats (Harlan Iberica, Barcelona, Spain) essentially as previously described (Aidil-Carvalho et al., 2017). Female rats were not used due to hormonal influences on LTP. Rats were anesthetized with halothane, decapitated, and both hippocampi were extracted in ice-cold artificial cerebrospinal fluid (aCSF), composition in mM: NaCl 124, KCl 3, NaH_2_PO_4_ 1.25, NaHCO_3_ 26, MgSO_4_ 1, CaCl_2_ 2, glucose 10, and gassed with a 95% O_2_ - 5% CO_2_ mixture.

### Electrophysiological recordings and induction of epileptiform activity

Hippocampal slices (Aidil-Carvalho et al., 2017) were kept in a resting chamber in the same gassed aCSF at room temperature 22ºC–25ºC for at least 1 h to allow their energetic and functional recovery, then one slice at a time was transferred to a submerged recording chamber of 1 ml capacity, where it was continuously superfused at a rate of 3 ml/min with the same gassed solution kept at 32ºC. Stimulation (rectangular pulses of 0.1 ms) was delivered through a bipolar concentric wire electrode placed on the Schaffer collateral/commissural fibres in the *stratum radiatum.* Two separate sets of the Schaffer collaterals (S1 and S2) were stimulated alternately every 10 s, each pathway being stimulated every 20 s (0.05Hz, Fig. 1A). Extracellular recordings of field excitatory post-synaptic potentials (fEPSP) were obtained from CA1 *stratum radiatum* through micropipettes 4M NaCl filled (2–4 MΩ resistance). fEPSPs were evoked on the two pathways, and the initial intensity of the stimulus, of comparable magnitude in both pathways, elicited a fEPSP of 600–1000 mV amplitude (near 50% of the maximal response), yet avoiding signal contamination by the population spike. The averages of six consecutive evoked fEPSPs from each pathway were obtained, measured, graphically plotted and recorded for further analysis with a personal computer using the LTP software (Anderson and Collingridge, 2001). The fEPSPs were quantified as the slope of the initial phase of the potential.

**Figure 1.**
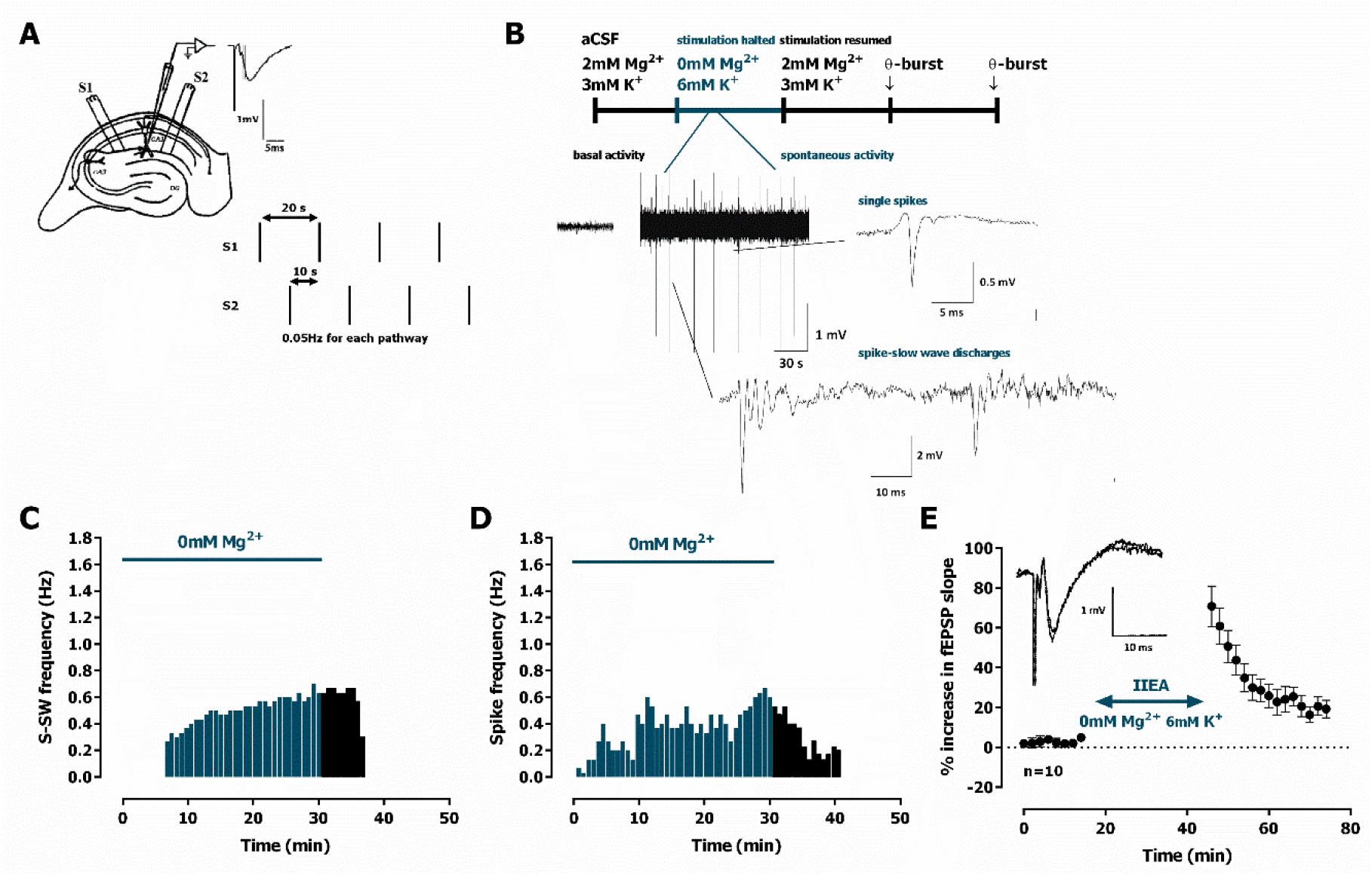
Interictal-like epileptiform activity induced by *0Mg^2+^* and LTP-like effect. **A.** Electrophysiological recordings of fEPSPs evoked by electrical stimulation in the CA1 area of hippocampal slices. The same recording configuration was used when monitoring spontaneous activity, but stimulation was halted during the 30-min perfusion with aCSF lacking Mg^2+^ and containing 6mM K^+^ *(**0Mg^2+^**).* **B.** Time-course of a typical experiment showing the spontaneous activity recorded prior to ***0Mg^2+^*** arrival to the superfusion chamber (left) and during the 30-min ***0Mg^2+^*** perfusion. Traces of the two main types of epileptiform activity recorded, single ***population spikes*** and ***spike-slow wave*** activity, are shown at the millisecond scale. **C-D.** Time course of the frequency of these two main types of spontaneous activity obtained during a single experiment. **E.** Averaged time-course of changes in fEPSP slope caused by ***0Mg^2+^*** perfusion. ***Inset***: traces of fEPSPs obtained in the same experiment before and 25-30 min after ***0Mg^2+^*** perfusion Traces are the average of eight consecutive responses and are composed of the stimulus artifact, the presynaptic volley and the fEPSP. Individual values (**C.** - **D.**) and the mean ± S.E.M are depicted (**E.**).

Spontaneous epileptiform activity (EA) was induced after a stable fEPSP slope baseline was obtained for at least 20min. Electrical stimulation was halted during these EA induction protocols to allow for the monitoring of pure spontaneous activity. Interictal-like epileptiform activity was induced by supressing Mg^2+^ in the superfusion media for 30 min (***0Mg^2+^***) while increasing the K^+^ concentration to 6mM. These concentrations were chosen to assure the short-term onset of EA (Gloveli et al., 1995; Antonio et al., 2016). Ictal-like EA was induced by perfusion with aCSF containing bicuculline (10 μM, ***Bic***) for 15 min. Electrical stimulation was resumed when spontaneous activity halted.

### LTP experiments

LTP was induced by a mild theta burst stimulation (TBS) pattern (five trains of 100 Hz, 4 stimuli, separated by 200 ms). The stimulation protocol used to induce LTP was delivered 30, 60 and 90 min after termination of EA induction protocols, provided a stable baseline was obtained for at least 16 min. LTP in control conditions (absence of prior EA) was time-matched to experiments performed 30 min after EA. Stimulus intensity was not altered during these stimulation protocols. LTP intensity was calculated as the % change in the average slope of the potentials taken from 50 to 60 min after TBS, compared to the average slope of the fEPSP measured during the 12 min that preceded TBS. Control and test conditions (for which epileptiform activity was induced prior to TBS) were tested in independent slices. LTP induced at different time points following EA was in some (but not all) cases tested in independent pathways on the same slice. In all experiments S1 always refers to the first pathway (left or right, randomly assigned) to which TBS was applied. Each *n* represents a single LTP experiment) performed in one slice from an independent animal, i.e., *n* denotes the number of animals.

The independence of the two pathways regarding LTP expression was tested at the end of the experiments by studying paired-pulse facilitation (PPF) across both pathways, less than 10% facilitation being usually observed. To elicit PPF, the two Schaffer pathways were stimulated with 50 ms interpulse interval. The synaptic facilitation was quantified as the ratio P2/P1 between the slopes of the fEPSP elicited by the second P2 and the first P1 stimuli.

### Western blot analysis of synaptic lipid raft markers and AMPA receptor changes influencing synaptic plasticity., GluA1 phosphorylation

Hippocampal slices were prepared as described above and allowed for functional recovery. Then 4 slices were introduced in placed in 100 μl Perspex chambers and superfused with gassed aCSF at 32 ºC at a flow rate of 3 ml/min. Field stimulation was delivered once every 15s in the form of rectangular pulses (1 ms duration, and an amplitude of 8 V) through platinum electrodes located above and below the slices. The electrical pulses were continuously monitored with an oscilloscope. In test (but not in control) slices interictal-like and ictal EA was induced with ***0Mg^2+^*** and ***Bic*** perfusion as described above for electrophysiological experiments. Stimulation lasted for 30 min before induction of EA, it was halted during EA and was resumed for 30 min (average duration of an electrophysiological experiment until first TBS delivery), after which slices were collected. Hippocampal synaptosomes were isolated from hippocampal slices as previously described (Cunha-Reis et al., 2017). Briefly, at the end of stimulation, slices were collected in sucrose solution (320mM Sucrose, 1mg/ml BSA, 10mM HEPES e 1mM EDTA, pH 7,4) containing protease (complete, mini, EDTA-free Protease Inhibitor Cocktail, Sigma) and phosphatase (1 mM PMSF, 2 mM Na3VO4, and 10 mM NaF) inhibitors and homogenized with a Potter-Elvejham apparatus. Each sample (n=1) was obtained from several slices (minimum 4 per condition) from 1-2 animals. The suspension was centrifuged at 3000g for 10 min, the supernatant collected and further centrifuged at 14000g for 12 min, and the pellet resuspended in 3 ml of a Percoll 45% (v/v) in modified aCSF (20mM HEPES, 1mM MgCl_2_, 1.2mM NaH_2_PO_4_, 2.7mM NaCl; 3mM KCl, 1.2mM CaCl_2_, 10mM glucose, pH 7.4). After centrifugation at 14 000 g for 2 min at 4ºC, the top layer (synaptosome fraction) was washed twice with modified aCSF also containing protease and phosphatase inhibitors and hippocampal membranes were resuspended at a concentration of 1mg/ml protein concentration (Bradford assay) in modified aCSF. Aliquots of this suspension of hippocampal membranes were snap-frozen in liquid nitrogen and stored at −80ºC until Western-blot analysis. All samples were analysed in duplicate in western-blot experiments.

For western blot, samples were incubated for 5 min at 95ºC with Laemmli buffer (125mM Tris-BASE, 4% SDS, 50% glycerol, 0,02% Bromophenol Blue, 10% β-mercaptoethanol), run on standard 10% sodium dodecyl sulphate polyacrylamide gel electrophoresis (SDS-PAGE) and transferred to PVDF membranes (Immobilon-P transfer membrane PVDF, pore size 0.45 μm, Immobilon) and blocked for 1 h with either 3% BSA or 5% milk (Caulino-Rocha et al., 2022). Membranes were incubated overnight at 4ºC with mouse anti-gephyrin (#147011, Synaptic Systems, AB_2810214), rabbit anti-PSD-95 (#CST-2507, Cell Signalling Tech., AB_561221), mouse anti-caveolin-1 (#ab106642, Abcam, AB_10861399), mouse anti-flotillin-1 (#ab133497, Abcam, AB _11156367), rabbit antiphospho-Ser845-GluA1 (1:2500, Abcam #Ab76321; RRID: AB_1523688), rabbit antiphospho-Ser-831-GluA1 (1:2000, Abcam #Ab109464; RRID: AB_10862154), rabbit anti-GluA1 (1:4000, Millipore # AB1504; RRID:AB_2113602), rabbit anti-GluA2 (1:1000, Proteintech #11994-1-AP; RRID: AB_2113725) and rabbit anti-alpha-tubulin (1:5000, Proteintech #11224-1-AP; RRID: AB_2210206) or rabbit anti-beta-actin (1:10000, Proteintech, Cat# 60008-1; RRID:AB_2289225) primary antibodies. After washing the membranes were incubated for 1h with anti-rabbit or anti-mouse IgG secondary antibody both conjugated with horseradish peroxidase (HRP) (Proteintech) at room temperature. HRP activity was visualized by enhanced chemiluminescence with ECL Plus Western Blotting Detection System using a ImageQuant™ LAS imager (GE Healthcare). Band intensity was estimated using the Image J software. ß-actin or α-tubulin band density was taken as a loading control and used for data normalization. The % phosphorylation for each AMPA GluA1 subunit target was calculated normalizing the change in phosphorylated form band intensity by the change in band intensity of the total GluA1 immunostaining.

## Materials

Bicuculline methochloride (Ascent Scientific, UK) was made up in 10mM stock solutions in DMSO. Aliquots of the stock solutions were kept frozen at −20ºC until use. An aliquot was thawed and diluted in aCSF for use in each experiment. The maximal DMSO concentration used was devoid of effects on fEPSP slope.

## Data and statistical analysis

LTP values are depicted as the mean ± S.E.M of *n* experiments. Each *n* represents a single experiment performed in slices obtained from one different animal for LTP experiments and a sample obtained from slices obtained from 1-2 animals for western blot experiments. Statistical analysis was performed using GraphPad Prism 6.01 for Windows. Significance of the differences between the LTP means was evaluated using paired Student’s t-test or repeated measures ANOVA with Sidak post-hoc test Repeated measures ANOVA with Sidak’s post-hoc test (when F was significant) was used to evaluate group differences in western blot experiments. No outliers were identified in our data (ROUT method).

## Results

The evoked fEPSPs recorded under basal stimulation conditions in the *stratum radiatum* of the CA1 area in hippocampal slices from young-adult rats (Fig. 1.A) had an average slope of 0.635±0.026 mV/ms (n=26), that represented 40-60% of the maximal response in each slice. Upon superfusion with modified aCSF lacking Mg^2+^ (***0Mg^2+^***) for 30min in the absence of electrical stimulation spontaneous activity started gradually with single spontaneous population spikes (Fig. 1.B, 0.7 to 1.2 mV amplitude, n=5) and 6-8 min later larger amplitude (2.7 to 4 mV, n=5) population spikes with wave reverberation (spike-slow wave discharges, Fig. 1.B) were observed (Fig. 1.C-D). All spontaneous activity was terminated within 8-12 min post ***0Mg^2+^*** washout. When resuming electrical stimulation an LTP-like effect was observed in the slope of the evoked fEPSPs (Fig. 1.E) with a potentiation of 21.0±4.7% (n=10) 30 min after resuming electrical stimulation.

When mild TBS was applied in control conditions the LTP magnitude corresponded to a 32.0±1.3% enhancement of fEPSP slope 50-60 min after TBS (n=13, Fig. 2.A and 4.A), an effect that was absent when slices were stimulated in the presence of the NMDA receptor antagonist AP-5 (100μM), as previously described (Rodrigues et al., 2021). When a TBS train was delivered 30 min past interictal-like EA induced by ***0Mg^2+^*** LTP expression was impaired (% increase in fEPSP slope: 18.0±1.8%, n=5, Fig. 2.B). When delivered to the slices 60 min past interictal-like EA, the same TBS stimulation elicited a larger LTP, nearly as the one obtained in control slices, now increasing by 24.8±1.1% (n=5) the fEPSP slope 50-60 min post TBS (Fig 2.C). This suggests interictal-like EA induced by ***0Mg^2+^*** does not promote marked or sustained changes in the LTP expressing ability of CA3 to CA1 synapses, that only appear to be transiently affected by this type of EA.

**Figure 2.**
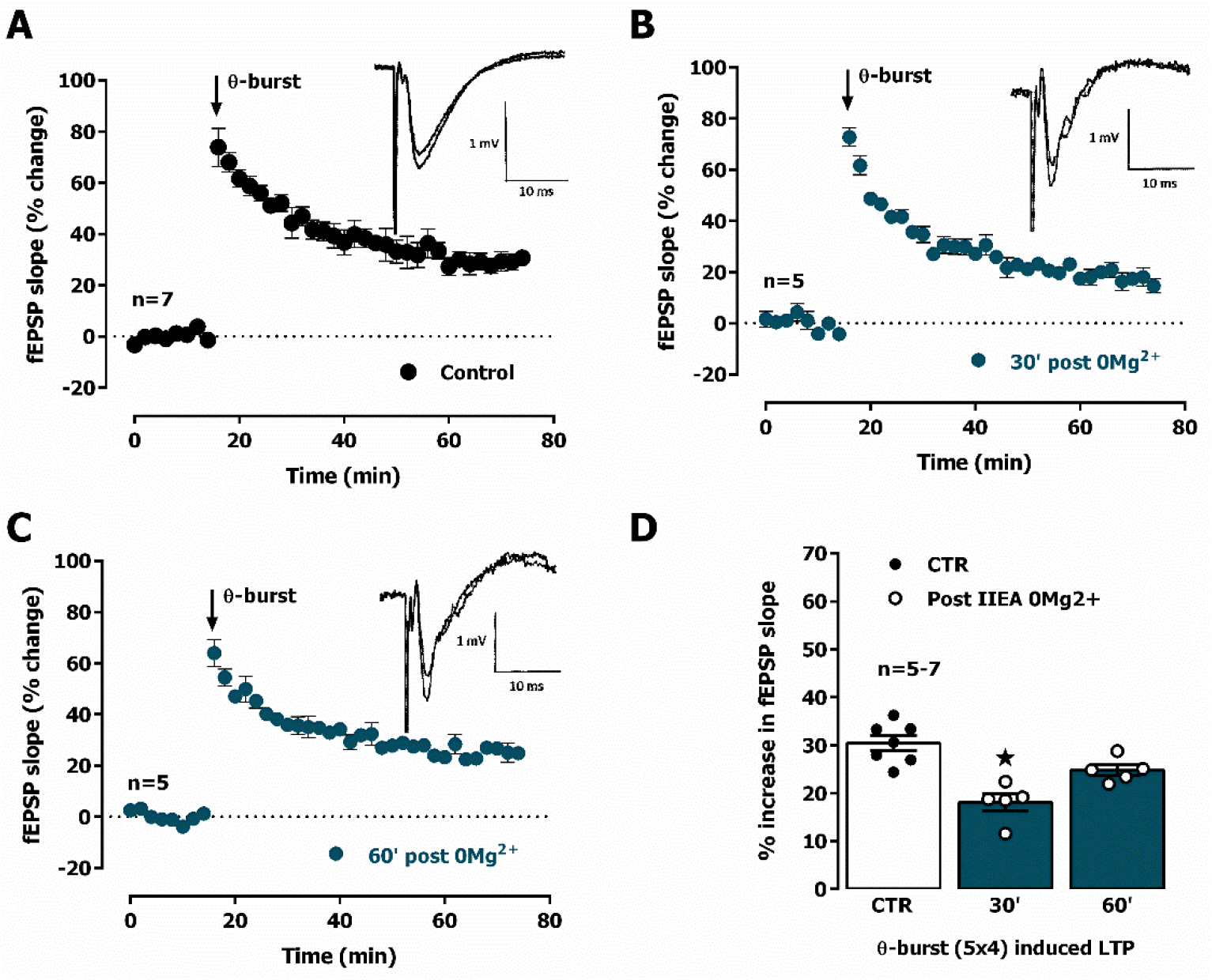
Interictal-like activity transiently impairs the expression of hippocampal CA1 long-term potentiation (LTP) of synaptic transmission. **A.** Averaged time-course of changes in fEPSP slope (-•-) caused by theta-burst stimulation (TBS, 5 bursts at 5 Hz, each composed of four pulses at 100 Hz) in experiments in which a control slice was stimulated in the absence of added drugs. ***Inset:*** Traces of fEPSPs obtained in one of the same experiments before and 50-60 min after TBS. Traces are the average of eight consecutive responses and are composed of the stimulus artifact, the presynaptic volley and the fEPSP. **B.** and **C.** Averaged time-course of changes in fEPSP slope caused by TBS applied either 30 min (**B.**) or 60 min (**C.**) after interictal epileptiform activity induced by ***0Mg^2+^*** perfusion (-•-) **D.** LTP magnitude estimated from the averaged enhancement of fEPSP slope observed 50-60 min after TBS in control slices (•, left open bar) or in slices previously experiencing interictal epileptiform activity induced by ***0Mg^2+^*** (○, blue bars). LTP was induced either 30 min (middle bar) or 60 min after epileptiform activity (right bar). ***Inset:*** Traces of fEPSPs obtained before and 50-60 min after TBS. Individual values and the mean ± S.E.M are depicted (**D.**). *p < 0.05 (Student’s t test) as compared to LTP magnitude in control slices (○, in the left).

Upon superfusion with modified aCSF containing bicuculline (10 μM, ***Bic***) for 16 min spontaneous activity started with single isolated population spikes (Fig. 3.A, 0.51 to 0.89 mV amplitude, n=7) that were increasingly more frequent than the ones observed during ***0Mg^2+^*** perfusion (Fig. 3.C). Within 4-6 min of ***Bic*** superfusion slower waves of synchronous activity (0.29 to 0.68 mV, n=7, Fig. 3.C) began to develop showing occasional high amplitude bursts (EPSP / slow-wave discharges, Fig. 3.A, and C). All spontaneous activity was terminated within 16-20 min post ***Bic*** washout. When resuming electrical stimulation an LTP-like effect was observed in the slope of the evoked fEPSPs (Fig. 3.D) with a potentiation of 44.8±8.7% (n=9) 30 min after resuming electrical stimulation.

**Figure 3.**
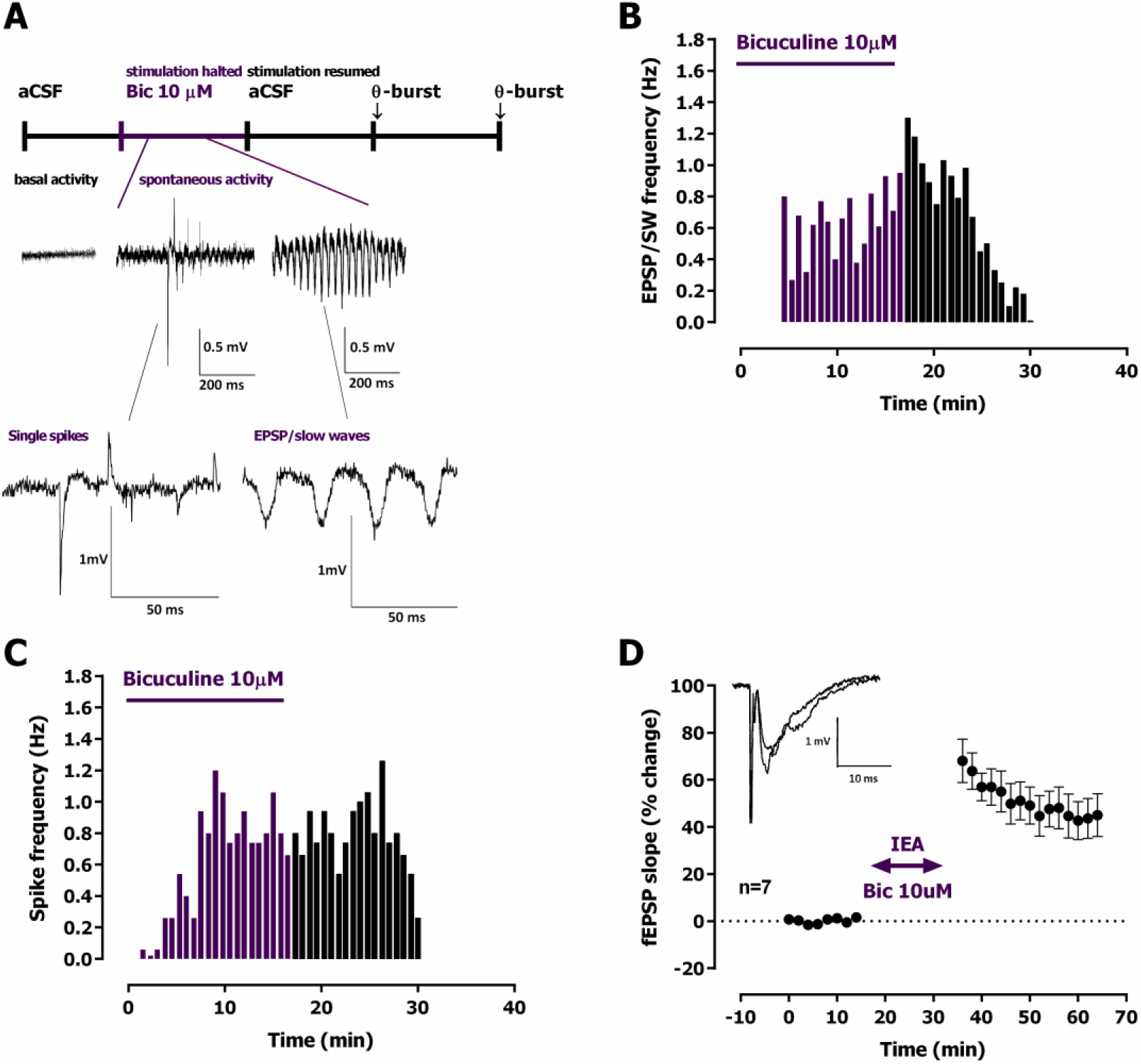
Ictal-like epileptiform activity induced by *Bicuculine* and LTP-like effect. **A.** Time-course of a typical experiment showing the spontaneous activity recorded prior to bicuculline (10 μM, ***Bic***) delivery to the superfusion chamber (left) and during the 16-min ***Bic*** perfusion. Specific examples of the two main types of epileptiform activity recorded, single ***population spikes*** and ***EPSP-slow wave*** activity, are shown at the millisecond scale. Stimulation was halted during ***Bic*** perfusion. **B-C.-** Time course of the frequency of these two main types of spontaneous activity obtained during a single experiment. **D.** Averaged time-course of changes in fEPSP slope caused by ***Bic*** perfusion. ***Inset***: traces of fEPSPs obtained in the same experiment before and 25-30 min after ***Bic*** perfusion. Traces are the average of eight consecutive responses and are composed of the stimulus artifact, the presynaptic volley and the fEPSP. Electrophysiological recordings of fEPSPs evoked by electrical stimulation were performed in the CA1 area of hippocampal slices. Individual values (**B.** - **C.**) and the mean ± S.E.M are depicted (**D.**).

LTP expression 30 min following ictal-like EA induced by ***Bic*** was strongly impaired (% increase in fEPSP slope: 12.5±2.2%, n=5, Fig. 4.B). TBS stimulation delivered to the slices 60 min past ictal-like EA, still elicited an impaired LTP, i.e., smaller than the one obtained in control slices, now increasing by 22.2±1.9% (n=5) the fEPSP slope 50-60 min post TBS (Fig 4.C). When delivered to the slices 90 min past ictal-like EA, TBS stimulation now induced an LTP larger than the one obtained in control slices, thus increasing by 45.0±3.0% (n=5) the fEPSP slope 50-60 min post TBS (Fig 4.D). As such, a biphasic change in the ability of TBS to induce LTP at CA1 hippocampal synapses occurs, first reflecting an impairment, then an enhanced capacity to elicit LTP following ictal-like EA induced by TBS.

**Figure 4.**
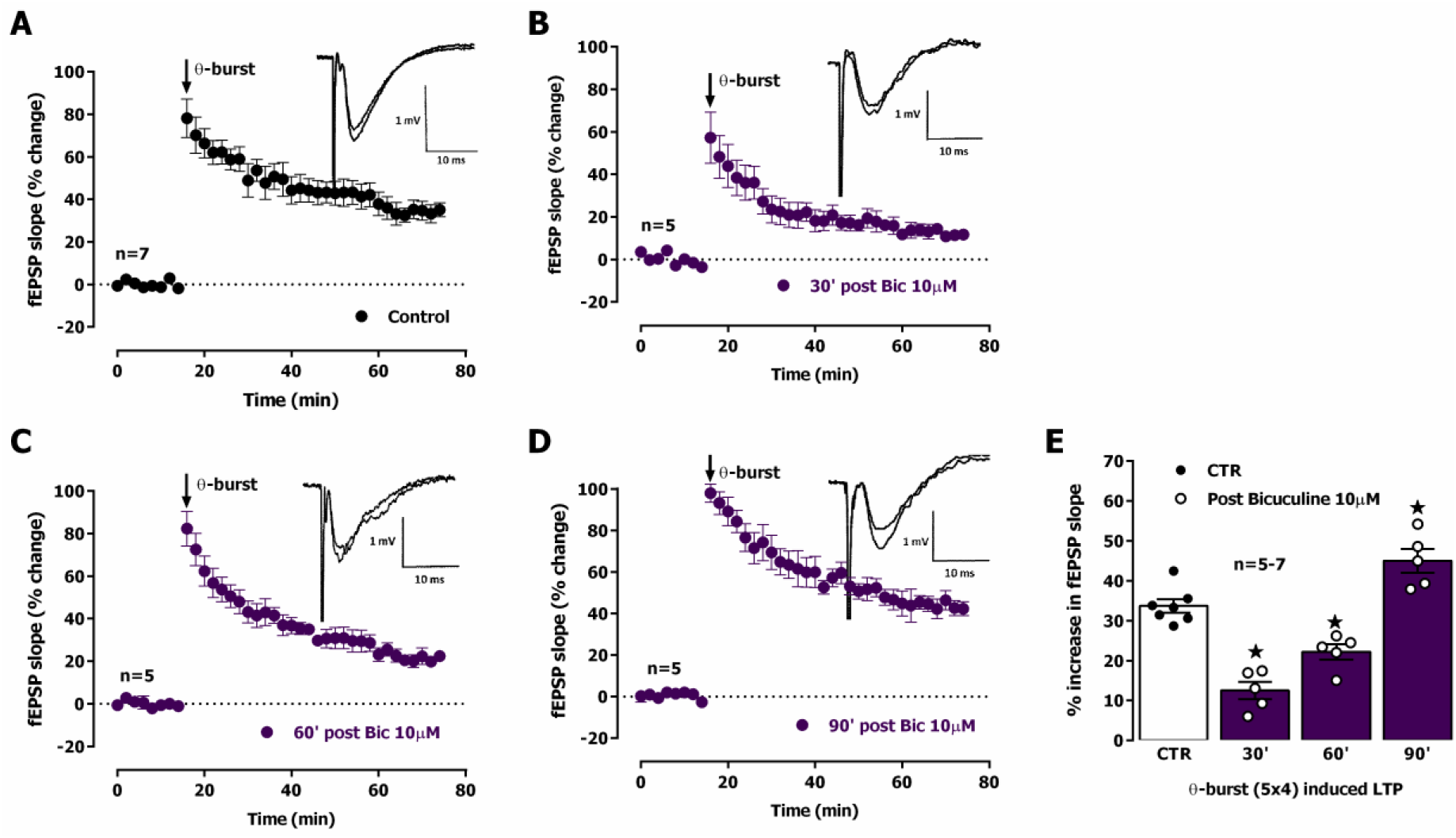
Ictal-like activity induces a biphasic alteration in the expression of hippocampal CA1 long-term potentiation (LTP) of synaptic transmission. **A.** Averaged time-course of changes in fEPSP slope (-•-) caused by theta-burst stimulation (TBS, 5 bursts at 5 Hz, each composed of four pulses at 100 Hz) in experiments in which a control slice was stimulated in the absence of added drugs. ***Inset:*** Traces of fEPSPs obtained in one of the same experiments before and 50-60 min after TBS. Traces are the average of eight consecutive responses and are composed of the stimulus artifact, the presynaptic volley and the fEPSP. **B.** - **D.** Averaged time-course of changes in fEPSP slope caused by TBS delivered either 30 min (**B.**), 60 min (**C.**) or 90 min (**D.**) after ictal epileptiform activity induced by bicuculline 10 μM *(**Bic**)* perfusion (-•-) **E.** LTP magnitude estimated from the averaged enhancement of fEPSP slope observed 50-60 min after TBS in control slices (•, left open bar) or in slices previously experiencing interictal epileptiform activity induced by ***Bic*** (○, purple bar). LTP was induced either 30 min (second bar), 60 min (third bar) or 90 min (last bar) after epileptiform activity. ***Inset:*** Traces of fEPSPs obtained before and 50-60 min after TBS. The mean ± S.E.M is depicted (**A.** - **D.**). *p < 0.05 (Student’s t test) as compared to LTP magnitude in control slices (○, in the left).

To further understand the molecular alterations at synapses that were underlying these changes in LTP expression we performed parallel experiments where hippocampal slices were also subjected to EA induced by ***0Mg^2+^*** or ***Bic***, and 30 min later were processed for synaptosome isolation and analysis of synaptic proteins associated with altered synaptic plasticity. These included AMPA receptor subunits GluA1 and GluA2, GluA1 phosphorylation in Ser831 and Ser845, Kv4.2 potassium channels, synaptic structural and lipid raft markers such as PSD-95, gephyrin, caveolin-1, flotillin-1, α-tubulin, and synaptophysin-1.

AMPA receptor subunit composition (GluA1-4) influences channel function and LTP outcomes (Chater and Goda, 2022), since AMPA receptors lacking GluA2 are Ca^2+^ permeable and when activated can further add to the postsynaptic Ca^2+^ rise and LTP levels. To elucidate the possible changes in synaptic AMPA receptor subunit composition that may impact its Ca^2+^ permeability and its contribution to altered LTP expression following EA we investigated changes in synaptic GluA1 and GluA2 levels and GluA1/GluA2 ratio. Following EA, the synaptic levels of AMPA GluA1 subunits (Fig. 5.A) were decreased to 77.1±9.0% (n=5) after ***0Mg^2+^*** exposure and to 72.0±11.8% (n=5) after ***Bic*** exposure. Opposingly, synaptic GluA2 subunits (Fig. 5.D) were increased to 127.3±10.1% (n=5) after ***0Mg^2+^*** exposure and to 152.8±14.3% (n=5) after ***Bic*** exposure. Consequently, GluA1/GluA2 ratios (Fig 5.E) were also markedly reduced after ***0Mg^2+^*** and ***Bic*** exposure. No significant differences were found between ***0Mg^2+^*** or ***Bic*** exposure for both targets as well as GluA1/GluA2 ratios.

**Figure 5.**
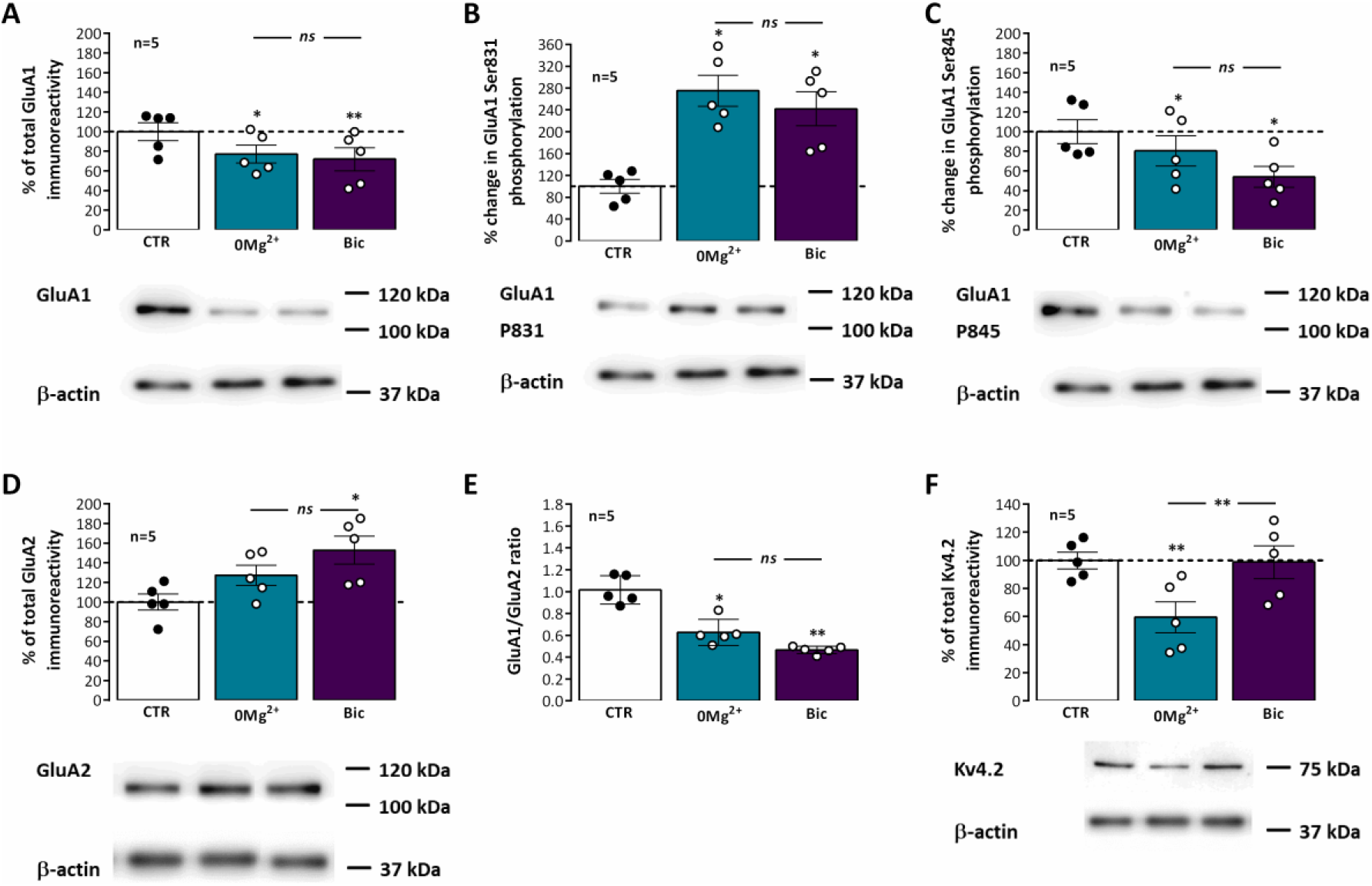
Impact of epileptiform activity on phosphorylation of synaptic hippocampal AMPA GluA1 subunits on Ser 831 and Ser 845, GluA1/GluA2 ratio and Kv4.2 levels. Each panel shows at the bottom the western-blot immunodetection of AMPA total GluA1 subunits (**A.**), its phosphorylated forms in Ser831 (**B.**) and Ser845 (**C.**), total GluA2 subunits (**D.**) and Kv4.2 potassium channels (**F.**) obtained in one individual experiment where hippocampal slices underwent Schaffer collateral basal stimulation for 20 min and then were subjected either to interictal-like epileptiform activity (EA) induced by perfusion with ***0Mg^2+^*** for 30 min or ictal-like EA by exposure to bicuculine (10 μM, ***Bic***) for 16 min, and then allowed to recover for 30 min before slice collection. Control slices were monitored for 70 min (the equivalent time of the EA protocol) before WB analysis. Western blot experiments were performed using synaptosome preparations obtained from these slices. Respective total GluA1 immunoreactivity (**A.**), % GluA1 phosphorylation in Ser831 (**B.**) or Ser845 (**C.**), residues, total GluA2 immunoreactivity (**D.**), GluA1/GluA2 ratios (**E.**) and total Kv4.2 potassium channel levels (**F.**) are also plotted at the top for each panel. Individual values and the mean ± S.E.M of five independent experiments are depicted. 100% - averaged GluA1 or GluA2 immunoreactivity or GluA1 phosphorylation obtained in control conditions (CTR, absence of EA). **\*** P < 0.05 *(ANOVA,* Sidak’s multiple comparison test) as compared to CTR; ***ns*** represents non -significant differences P > 0.05 (*ANOVA*, Sidak’s multiple comparison test) between respective bars.

LTP expression, and particularly mild TBS induced LTP, depends on the Ca^2+^-dependent auto-phosphorylation of Ca^2+^/calmodulin dependent protein kinase II (CaMKII), that influences the phosphorylation of AMPA GluA1 subunits and their synaptic recruitment (Appleby et al., 2011; Rodrigues et al., 2021). These are also targeted by other intracellular kinases like protein kinase A and C (PKA and PKC) affecting hippocampal LTP expression by supporting traffic or altering AMPA receptor opening probability (Lee et al., 2003). Although not directly involved in mild TBS-induced LTP (Rodrigues et al., 2021), altered PKA and PKC activity during EA could influence AMPA basal phosphorylation thus influencing subsequent TBS-induced LTP. We thus investigated if altered GluA1 phosphorylation levels at Ser831, a prominent target of CaMKII and PKC, and Ser845, a main target of PKA were implicated in altered LTP responses following EA. Phosphorylation of GluA1 subunits in Ser831 (Fig 5.B) was markedly increased to 275.0±12.7% (n=5) after ***0Mg^2+^*** induced EA and 241.9±31.1% (n=5) after ***Bic*** induced EA. Opposingly, phosphorylation of GluA1 subunits in Ser845 (Fig 5.C) was decreased to 80.4±15.4% (n=5) after ***0Mg^2+^*** exposure and 53.9±10.6% (n=5) after ***Bic*** exposure. No significant differences were found between ***0Mg^2+^*** or ***Bic*** exposure for the phosphorylation on both targets.

Kv4.2 K^+^ channels are important regulators of dendritic excitability by mediating the transient A-current. Its membrane levels and phosphorylation status are regulated by electrical activity patterns used in LTP induction (e.g. strong TBS), suggesting that the activity of the channel, contributes to LTP expression (Frick et al., 2004; Kim and Hoffman, 2008; Rodrigues et al., 2021). Following EA, Kv4.2 levels (Fig. 5.F) were reduced to 59.5±11.1% (n=5) after ***0Mg^2+^*** exposure but were not significantly altered (98.8±11.7%, n=5) after ***Bic*** induced EA.

Synaptic lipid rafts are membrane microdomains that can be associated with classic lipid raft markers like caveolin-1 and flotillin-1, or that can be identified by synaptic lipid-raft nucleating/associated proteins like PSD-95 or gephyrin. Following EA, the synaptic levels of flotillin-1 (Fig. 6.A) were markedly reduced to 34.3±4.1% (n=5) after ***0Mg^2+^*** exposure and 46.3±11.4% (n=5) after ***Bic*** exposure while the levels of caveolin-1 (Fig. 6.B) were dramatically reduced to 10.6±4.7% (n=5) after ***0Mg^2+^*** exposure and 21.9±5.6% (n=5) after ***Bic*** exposure. Regarding the postsynaptic scaffolding protein of glutamatergic synapses PSD-95, after EA, it was observed an increase in PSD-95 levels to 178.6±11.5% (n=5) following ***0Mg^2+^*** exposure and to 180.8±24.9% (n=5) upon ***Bic*** exposure. Gephyrin was also significantly altered 30 min next to EA, increasing to 222.9±24.6% (n=5) upon ***0Mg^2+^*** exposure and 246.8±27.8% (n=5) upon ***Bic*** exposure. Only for PSD-95 was observed a significant difference in the EA response between ***0Mg^2+^*** and ***Bic*** exposure.

**Figure 6.**
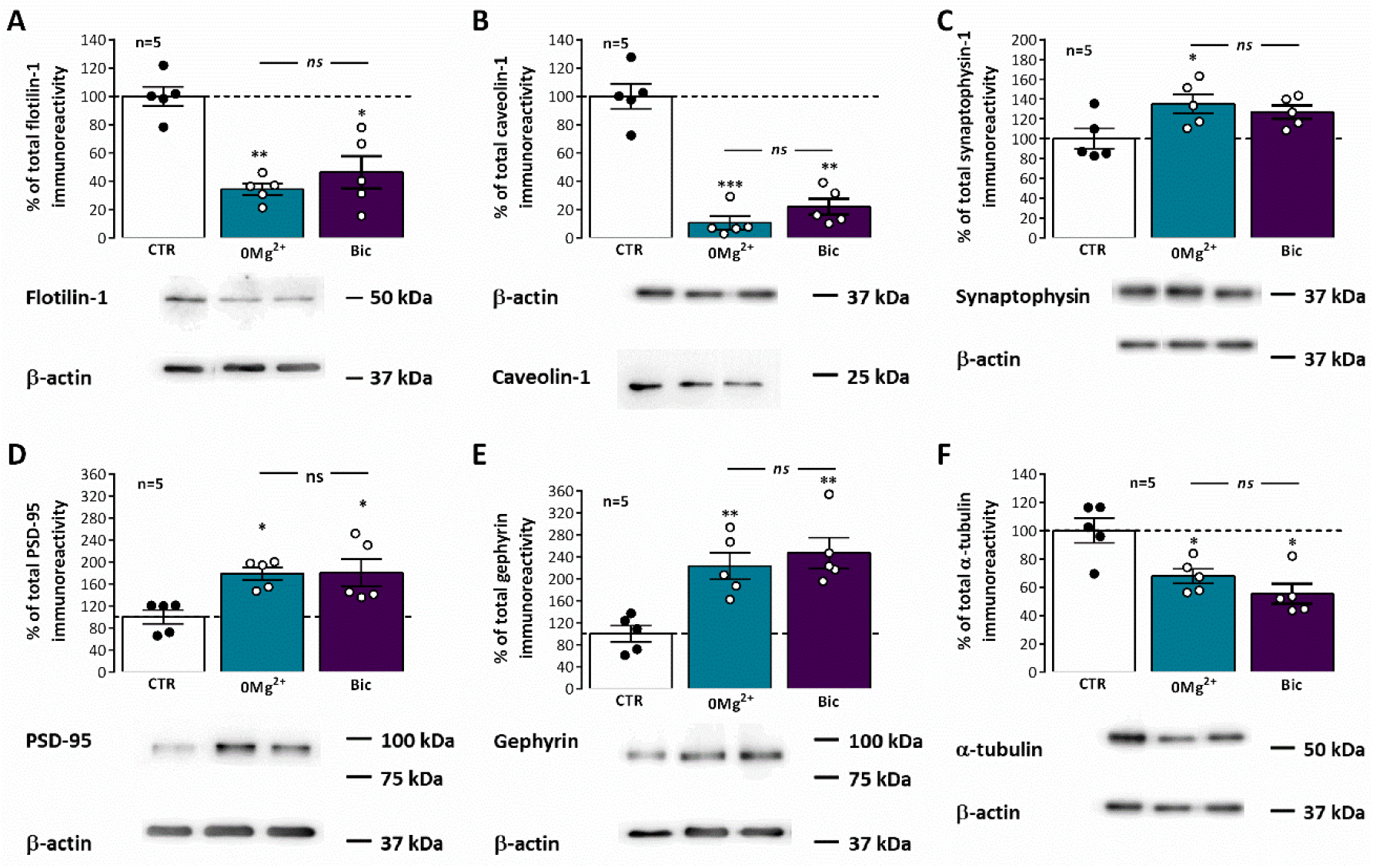
Impact of epileptiform activity on classic and synaptic lipid raft markers. Each panel shows at the bottom the western-blot immunodetection of flotilin-1 (**A.**), caveolin-1 (**B.**), synaptophysin-1 (**C.**) PSD-95 (**D.**), gephyrin (**E.**), and alpha-tubulin (**F.**) obtained in one individual experiment where hippocampal slices underwent Schaffer collateral basal stimulation for 20 min and then were subjected either to interictal-like epileptiform activity (EA) induced by perfusion with ***0Mg^2+^*** for 30 min or ictal-like EA by exposure to bicuculine (10 μM, ***Bic***) for 16 min, and then allowed to recover for 30 min before slice collection. Control slices were monitored for 70 min (the equivalent time of the EA protocol) before WB analysis. Western blot experiments were performed using synaptosome preparations obtained from these slices. Respective average change in total flotilin-1 (**A.**), caveolin-1 (**B.**), synaptophysin-1 (**C.**) PSD-95 (**D.**), gephyrin (**E.**), and alpha-tubulin (**F.**) immunoreactivities are also plotted at the top in each panel. Individual values and the mean ± S.E.M of five independent experiments are depicted. 100% - averaged PSD-95, gephyrin, caveolin-1, flotillin-1, or synaptophysin-1 immunoreactivity in control conditions (CTR, absence of EA). **\*** P < 0.05 *(ANOVA,* Sidak’s multiple comparison test) as compared to CTR; ***ns*** represents non -significant differences P > 0.05 (*ANOVA*, Sidak’s multiple comparison test) between respective bars.

To understand the relation between the changes in these different proteins at synapses and overall synaptic alterations in this sinaptossomal preparation we also evaluated the synaptic levels of synaptophysin-1, a synaptic vesicle associated glycoprotein. Synaptophysin-1 levels were mildly enhanced following exposure to both *0Mg^2+^* (135.1±10.0%, n=5) or *Bic* (126.9±6.7%, n=5). In addition, we investigated changes in the structural protein α-tubulin, since our preliminary studies using it as a western blot loading control suggested that the synaptic levels of this protein were also significantly changing after EA. The synaptic levels of α-tubulin were decreased to 67.7±5.1% (n=5) following *0Mg^2+^* exposure and 55.0±7.0% (n=5) after *Bic* exposure.

## Discussion

In the present work we first describe that: 1) ictal-like EA induces an impairment in TBS-induced LTP within 30 min −1h after the insult and this is reversed to a mild enhancement 1h 30m after; 2) interictal-like EA induces only a mild impairment in TBS-induced LTP observed 30 min after the insult and this is fully recovered 1h after; 3) Impairment of LTP induced by EA was globally associated to a decrease in AMPA GluA1 subunit levels and an increase in GluA2 subunit levels, resulting in a marked decrease of the GluA1/GluA2 ratio, together with a marked increase in GluA1 P831 phosphorylation and a mild decrease in GluA1 P845 phosphorylation; 4) Kv4.2 K^+^ channels levels were decreased following interictal-like EA but not following ictal-like EA; 5) Impairment of LTP induced by EA induced a decrease in the levels of classic lipid raft markers caveolin-1 and flotillin-1, and an increase in synapse specific lipid raft markers PSD-95 and, most prominently, gephyrin.

The physiological alterations observed in this work for LTP induction and expression, as well as for the molecular composition of synapses provide a good insight on the very early alterations on synaptic plasticity in adult rodents following seizure-like activity (the one induced by bicuculine, 10μM) and following a pattern of neuronal interictal-like activity (induced by Mg^2+^ suppression), often observed in the latent period of epileptogenesis in animal models, and the characteristic brain activity of epilepsy patients between seizures, as its denomination suggests. The fact that LTP expression is influenced by previous neuronal and/or synaptic activity is not itself new (Abraham and Bear, 1996; Colgin et al., 2004; Zhang et al., 2005) and previous studies reported structural alterations in synapse architecture following seizures (Leite et al., 2005; Postnikova et al., 2017). Furthermore, altered synaptic plasticity was observed following numerous physiologic stress conditions, (Félix-Oliveira et al., 2014; Arias-Cavieres et al., 2021) that were also associated with clustering of synaptic markers and altered synaptic morphology (Sebastian et al., 2013). Nonetheless, the evaluation of LTP changes occurring following *in vitro* events resembling seizure-like activity versus the ones caused by altered resting neuronal activity observed in epileptic patients was never performed. The use of the hippocampal slice *in vitro* system for seizure generation was preferred to the *in vivo* model of epilepsy also used by our research group (Serpa et al., 2022), since it allowed us to assess LTP in the first hours following EA, thus avoiding the dissection and recovery time after slice preparation (usually 1h - 1h 30m) that would expand the time window of LTP inspection. This is of particular interest to the future development of early intervention therapies to prevent epileptogenesis.

Interictal-like epileptiform activity, lasting for about 30 min, was characterised by spaced large amplitude burst of neuronal activity separated by more silent periods with occasional single spikes while ictal like activity, lasting for 10-15 min, was characterised by a continuous activity showing both population spikes and EPSPs. As previously observed by many others (Gloveli et al., 1995; Debanne et al., 2006; Postnikova et al., 2020; Ergina et al., 2021), synaptic transmission was potentiated following both ictal-like and interictal-like EA (Fig. 1.E and Fig. 3.D), an LTP-like effect of EA similarly dependent on NMDA receptor activation (Ben-Ari, 2001; Debanne et al., 2006). In this paper, we also tested the ability of EA to condition further LTP induction by mild TBS (5 trains of 4 pulses delivered at 100Hz with 200ms interburst interval). We observed a transient decrease in LTP expression when induced 30 min following inter-ictal like EA (Fig. 2) compared to time-matched control slices that was virtually non-existing when LTP was induced 1h following that insult. However, the impairment in LTP expression was much stronger when LTP was induced 30 min following ictal-like activity (Fig. 4), and was still present, although to a minor extent, for LTP induced 1h following ictal-like activity. In addition, LTP induced 1h 30 min after ictal-like activity was slightly larger than the one observed in control conditions. Previous studies addressed the alterations in LTP following in vitro 4-AP induced ictal-like EA in hippocampal slices or 3h after Li^2+^-pilocarpine-induced seizures (Kryukov et al., 2016; Postnikova et al., 2020), yet these studies differ from this in that they used juvenile rodents (approx. 3-week-old). Both studies encountered a small reduction of LTP, partially agreeing with the observations in our study, but using a slightly stronger TBS pattern, likely a requirement to induce a robust LTP in juvenile rats, since CA1 LTP induced by TBS undergoes a post-weaning maturation from 3-12 weeks of age in this model (Rodrigues et al., 2021). Interestingly, we observed a more pronounced impairment in LTP following ictal-like activity and a near full recovery within 1h-1h 30m after EA. This may be due to the shorter duration of the ictal-like activity induced in this paper or may reflect differences in the *in vitro* EA induction methodologies (4-AP vs bicuculine). The differences in the age, the LTP and EA induction methods and/or in the time frame of LTP evaluation after EA preclude a more elaborated discussion of our work against these studies. However, another important aspect of our observations is that ictal-like activity, although shorter in duration than interictal-like activity, had a much greater impact on hippocampal LTP. This is in line with previous theories arguing that interictal-like activity is protective in epileptogenesis (Avoli, 2001; Avoli et al., 2006) and suggests that altered LTP mechanisms are not only important for epileptogenesis but also a relevant therapeutic target in its prevention (Albensi et al., 2004, 2007; Avanzini et al., 2014).

LTP induction in the CA1 area of the hippocampus classically relies on the postsynaptic Ca^2+^ rise generated by NMDA receptor activation (Bliss et al., 2018) that results from primary AMPA receptor mediated postsynaptic depolarization. AMPA receptor subunit composition, a variable tetrameric combination of GluA1-4 subunits influencing its channel function, is another factor influencing LTP outcomes (Wiltgen et al., 2010; Chater and Goda, 2022). AMPA receptors containing exclusively subunits GluA1,3 and 4 are Ca^2+^ permeable and when activated can further add to the postsynaptic Ca^2+^ rise and LTP levels, while GluA2 containing AMPA receptors are essentially impermeable to Ca^2+^. Synaptic AMPARs in the adult hippocampus are largely composed of GluA1–3 that form predominately GluA1/2 and GluA2/3 heterodimers, with GluA1/2 being the most common (Chater and Goda, 2022), yet diminished levels of GluA2 containing AMPA receptors were linked to aging and pathological conditions such as brain lesions. In this paper we found that the levels of GluA1 AMPA receptor subunits were decreased while GluA2 AMPA receptor subunit were increased in hippocampal synaptosomes obtained from hippocampal slices exposed to interictal-like and ictal-like activity (Fig 5.A and D). Consequently, GluA1/GluA2 ratio was increased for both conditions (Fig. 5.E). These changes were more accentuated following ictal-like activity. This suggests that epileptiform activity transiently reduces the number of Ca^2+^-permeable AMPA receptors at hippocampal synapses in adult rats, that is likely an endogenous neuroprotective measure. Interestingly, decreased Ca^2+^ permeability following ***0Mg^2+^*** induced EA was previously reported in mixed neuroglial cultures (Gaidin et al., 2019). Although this system has a very different circuit connectivity than hippocampal slices, and cultures are prepared from embryonic rats, this is consistent with our results reporting and increase in the GluA2 subunit component of AMPA receptors. Furthermore, recent works implicated synaptic GluA2-lacking Ca^2+^ permeable receptors in the LTP-like effects of 4-AP induced EA in juvenile rats (Postnikova et al., 2020), identifying such synaptic alterations as the main event altered following EA as opposed to other pyramidal neuron membrane properties (Ergina et al., 2021). Given the yet relative immaturity of hippocampal GABAergic circuits / GABAergic transmission and their contribution to ictogenesis and control of LTP at these very young ages (Papatheodoropoulos and Kostopoulos, 1998; Akgül et al., 2019; Rodrigues et al., 2021; Caulino-Rocha et al., 2022), the fact that GluA1-depedent mechanisms differentially contribute to TBS-induced LTP in juvenile and adult mice (Liu et al., 2020), and that diminished levels of GluA2 containing AMPA receptors were linked to postnatal developmental stages (Chater and Goda, 2022), it is likely that the AMPA glutamatergic component of this LTP-like effect may be distinct in adult circuitry (Goodkin et al., 2008; Avoli et al., 2022; Lévesque et al., 2022). The above-cited studies are nonetheless extremely relevant to understand early life epileptogenesis. Nonetheless, since the initiation of EA is itself dependent on GABAergic transmission (Avoli et al., 2022), the role of GABAergic transmission in the LTP-like effects of EA should also be elucidated. This would require a distinct approach to induce ictal-like activity than the one used in this study since direct effects of bicuculine on GABAergic receptors may alter also the long-lasting response to EA.

The experimental approach used in this paper to monitor changes in synaptic AMPA receptors strongly limits the detail obtained on their subsynaptic distribution (Chater and Goda, 2022). as it does not allow the distinction between synaptic and perisynaptic AMPA receptors, nor between pre- or post-synaptic receptors. In the context of seizure induced epileptogenesis, activation of extrasynaptic receptors by glutamate spillover from active synapses may be crucial to the establishment of metaplastic events contributing to maladaptive homeostatic plasticity mechanisms such as changes in AMPA/NMDA ratio at glutamatergic synapses (Postnikova et al., 2020; Ergina et al., 2021; Chater and Goda, 2022). As such, further elucidation of the exact changes occurring at synaptic and perisynaptic sites would be of great use in elucidating the contribution of AMPA receptors to early alterations in synaptic plasticity to epileptogenesis.

LTP expression depends on the Ca^2+^-dependent auto-phosphorylation of Ca^2+^/calmodulin dependent protein kinase II (CaMKII), that in turn influences the phosphorylation of AMPA GluA1 subunits and their synaptic recruitment (Appleby et al., 2011; Park et al., 2021). These are also targeted by intracellular kinases like protein kinase A and C (PKA and PKC) that modulate hippocampal LTP by supporting the traffic or altering the opening probability of AMPA receptors (Lee et al., 2003). Since epileptiform activity could be limiting hippocampal LTP by inducing an activity-dependent enhancement in the phosphorylation status of AMPA GluA1 subunits, thus generating a ceiling effect for LTP induction, we investigated the impact of epileptiform activity on the phosphorylation status of AMPA GluA1 receptors. The phosphorylation levels of AMPA GluA1 subunits at Ser831, a prominent target of CaMKII and PKC, was enhanced after both ictal-like and interictal like EA patterns, the effect being slightly more pronounced following interictal-like activity (Fig. 5.B). This suggests that this mechanism is also significantly contributing to the LTP-like effects of EA by increasing channel conductance and promoting the recruitment of AMPA receptors to the synapse (Derkach et al., 1999, 2007). This may also be the main cause for impaired LTP expression following EA activity, by limiting the additional phosphorylation at Ser831 that can be induced by TBS. Thus, although the overall number of GluA1 subunits is decreased, the remaining GluA1 containing AMPA receptors are more likely found at synapses. Our observations early following EA contrast with the observed decrease in GluA1 Ser831 phosphorylation observed 3h after pilocarpine induced seizures (Russo et al., 2013). Interestingly though, these results support our observations that a biphasic response to LTP induction occurs following seizures/EA, with decreased efficacy first observed and increased efficacy observed later.

The phosphorylation of GluA1 at Ser845, a main target of PKA, was contrariwise decreased following both types of EA, and this decrease was more pronounced following ictal-like activity (Fig. 5.C). This change opposes the one observed after common TBS protocols *in vitro* (Park et al., 2021) but is consistent with or findings of decreased GluA1 and GluA2 containing receptors at synapses. This may also reflect and endogenous neuroprotective measure against the consequences of hyperexcitability induced by EA, as phosphorylation of GluA1 subunits at Ser845 drives AMPA receptor synaptic insertion (Fig. 7), which would constitute a maladaptive homeostatic plasticity mechanism following seizures. This is fundamentally different from what is observed in the chronic period in the Li^2+^-pilocarpine model of epilepsy, where both GluA1 P831 and P845 phosphorylation are increased (Serpa et al., 2022).

**Figure 7.**
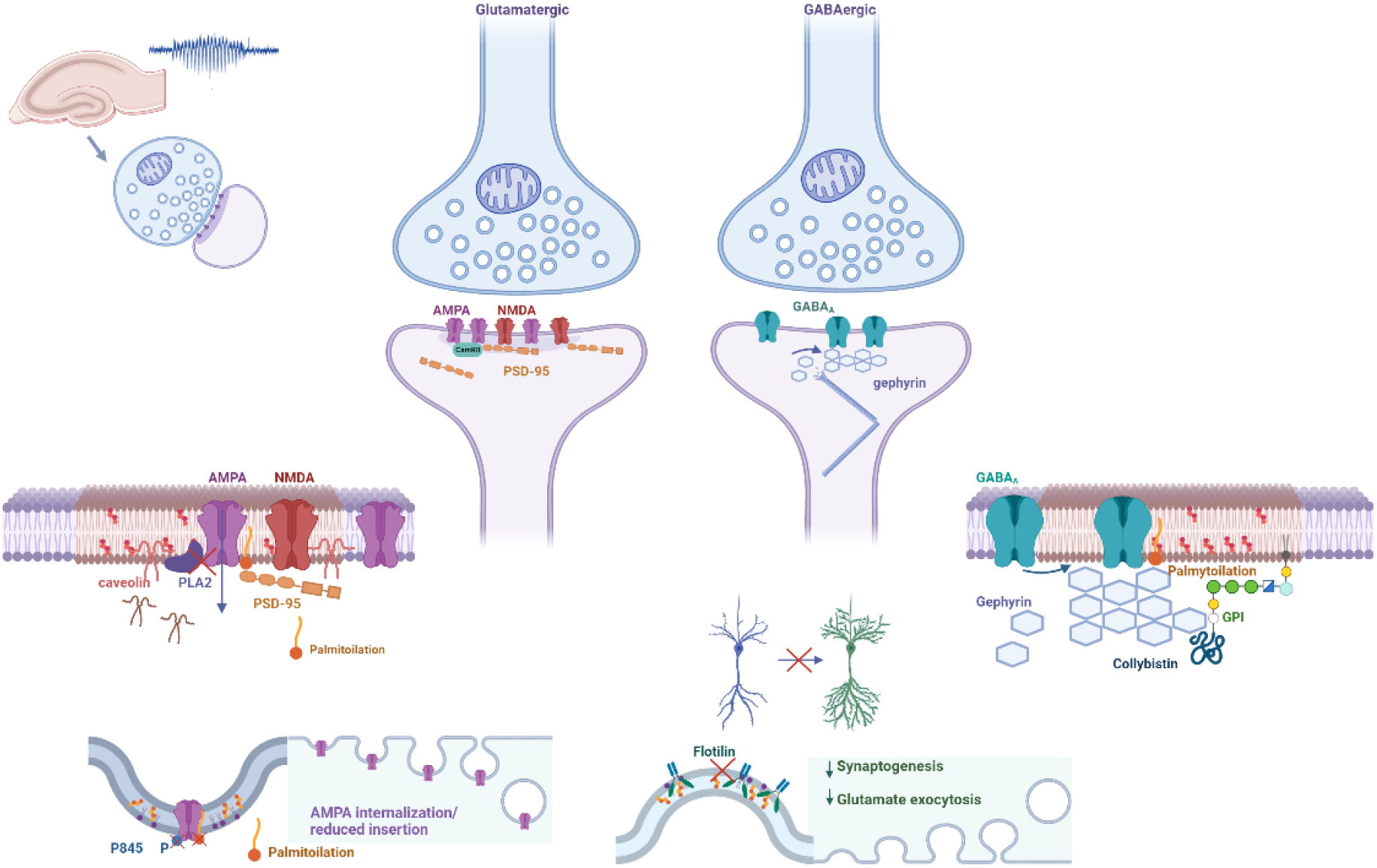
Proposed synaptic molecular mechanisms in altered synaptic plasticity and lipid raft dynamics following epileptiform activity. Altered LTP efficiency within the first hours of ictal-like and interictal-like EA may involve lipid raft dependent AMPA receptor internalization, impaired glutamatergic synaptogenesis, impaired GABA release and the GPI-dependent recruitment of GABAA receptors to GABAergic synapses.

In this work we also observed a decrease in synaptic Kv4.2 channels following interictal-like EA yet not following ictal-like EA (Fig. 5.F). The Kv4.2 channel is a K^+^ channel with more prominent expression in dendritic spines than in dendritic shafts, that is largely responsible for the Ca^2+^-activated delayed rectifying A-current (I_A_) in CA1 pyramidal neuron distal dendrites, where it acts to control signal propagation and compartmentalization (Kim et al., 2007; Beck and Yaari, 2008; Kim and Hoffman, 2008). LTP induction with strong TBS stimuli reduces Kv4.2 dendritic membrane levels, leading to a shift in the voltage-dependence of I_A_, which suggests that Kv4.2 channel activity contributes to LTP expression through modifications such as phosphorylation and/or trafficking (Frick et al., 2004; Kim and Hoffman, 2008). Kv4.2 channels are responsible for the precision of the time window allowed for LTP induction, since elimination of dendritic I_A_ in Kv4.2(-/-) mice results in an expanded time window, making LTP induction less dependent on the temporal relation of pre- and postsynaptic activity (Zhao et al., 2011). Interestingly, I_A_ activation can also influence synaptic NMDA receptor composition at CA1 pyramidal neurons (Jung et al., 2008) and is a determinant factor of synaptic morphology and seizure susceptibility (Tiwari et al., 2020). Altogether, this suggests that LTP induction will be easier to induce following interictal-like EA, resulting from an enlarged time window provided by Kv4.2 withdrawal from synapses. This may explain the lower impact and faster recovery of LTP expression following interictal-like vs ictal-like activity in our study. This effect is however contradictory with the overall observed putative neuroprotective synaptic molecular remodelling in AMPA receptor levels and phosphorylation.

In this paper we also found significant changes in α-tubulin levels in the sinaptossomal fraction obtained from hippocampal slices subjected to experimental conditions similar to the ones eliciting ictal-like and interictal-like EA (Fig. 6.F). As such, we could not use this protein as loading control for our western blot analysis. Although until recently the role of the microtubule cytoskeleton at synapses was not fully acknowledged recent evidence has shed light on its importance in synaptic function and evidence of its influence on neurotransmitter release and synaptic plasticity has been advanced (Waites et al., 2021). As such, decreased α-tubulin levels likely indicate structural changes in synaptic morphology that have previously been reported following epileptiform activity (Leite et al., 2005), suggesting that α-tubulin may undergo incorporation into microtubules following EA. Interestingly, the control of synaptic Ca^2+^ overload by microtubule dynamics has recently been reported (Vajente et al., 2019). Whether this mechanism may play a role endogenous neuroprotection against hyperexcitable states it remains to be established.

Chemical synapses are highly polarized subcellular structures comprising microdomains such as the presynaptic active zone and the postsynaptic density. In particular, the pre and post-synaptic membrane lipid composition is distinct from the remaining neuronal membrane, suggesting synaptic signalling is strongly dependent on this specialization (Borroni et al., 2016; Flores et al., 2019). This is based on the diversity of membrane lipid components that confers membranes with domains showing distinctive properties in terms of size, rigidity, and thickness (Wilson et al., 2020). Lipid rafts, liquid-ordered membrane nanodomains segregated from the bulk membrane and enriched in cholesterol, sphingolipids and GPI-anchored proteins, are important dynamic signalling platforms and comprise two main types of structure, caveolae, curved membrane domains, or planar lipid rafts (Allen et al., 2007; Borroni et al., 2016). These are characterized by the presence of the membrane proteins caveolins, a family of 22-kDa cholesterol-binding membrane protein, and flotilins, a family of 49-kDa membrane proteins associated with the membrane inner leaflet, respectively, that play an active role in promoting the phase separation within the membrane that maintains these synaptic lipid nanodomains (Allen et al., 2007; Frick et al., 2007). Nevertheless, lipid domain stability is also regulated by its lipid composition and lipid leaflet distribution (Sodero et al., 2011). Neurotransmitter release and neurotransmitter receptor dynamics relies on direct or indirect interactions of membrane proteins and receptors with these specific membrane domains (Allen et al., 2007; Borroni et al., 2016). Interestingly, upregulation of excitatory neurotransmission was shown to be a potent inducer of cholesterol loss, and this has been implicated as an endogenous neuroprotective mechanisms against oxidative stress (Iannilli et al., 2011; Sodero et al., 2011). Furthermore, the interaction of postsynaptic anchoring proteins like PSD-95 and gephyrin and neurotransmitter receptors, like AMPA receptor, with lipid raft domains is often regulated by synaptic activity through post-translational modifications like palmitoylation or GPI anchoring (Papadopoulos et al., 2015; Tulodziecka et al., 2016; Itoh et al., 2018). In this work we observed an extremely marked decrease in the synaptic levels of caveolin-1 and flotillin-1 (Fig. 6.A and B), suggesting that EA is a strong disruptor of normal lipid raft stability and of trafficking mechanisms regulated by these proteins. This may in turn affect enormously the synaptic signalling through AMPA, NMDA and GABA_A_ receptors and the synaptic plasticity mechanisms through them triggered. In fact, caveolin-1 enhances AMPA receptor binding capacity through direct inhibition of PLA2 activity, suggesting that a reduction of caveolin levels can impair LTP expression (Gaudreault et al., 2004). Furthermore, caveolin-1 is strongly involved in post-injury remodelling of the neuronal membrane (Gaudreault et al., 2005), while its deficiency has been implicated in enhanced cerebral ischemic injury (Jasmin et al., 2007), suggesting that caveolins are essential for neuronal membrane recovery following different insults. Flotilin-1 in turn is strongly implicated in glutamatergic synapse maturation, dendritic pruning and was reported to enhance glutamate release (Swanwick et al., 2010). In addition, although classically associated with planar lipid rafts, flotillin proteins were also identified as structural and functional components of a clathrin-independent endocytic pathway, inducing membrane curvature and the formation of plasma-membrane invaginations morphologically similar to caveolae (Frick et al., 2007), and were reported to be involved in neurotransmitter transporter internalization (Cremona et al., 2011). As such, decreased levels of flotillin-1 following seizures may account both for decreased synaptic stability, impaired glutamatergic transmission and contribute to impaired synaptic plasticity observed in this work. Alternatively, flotilins may be degraded intracellularly in an endogenous neuroprotective effort to reduce hyperexcitability. Altogether, our experimental observations suggest that caveolins and flotilins are promising therapeutic targets in epileptogenesis, but the exact mechanisms to be targeted still need further investigation.

One possible explanation for the massive loss of some synaptic lipid raft markers could be the occurrence of massive synaptic protein internalization, degradation and autophagy in response to synaptic activity during hyperexcitable states (Li et al., 2012; Soykan et al., 2021). This would imply a reduction in nerve terminal synaptic vesicles and its characteristic markers. Our observations that the synaptic vesicle marker synaptophysin is not significantly altered following both ictal-like and interictal-like EA (Fig. 6.C), suggests that these mechanisms are not likely to contribute to the observed reduced levels of synaptic proteins in this work.

In contrast with the levels of classic lipid raft markers, we also observed that the synaptic levels of postsynaptic scaffolding protein PSD-95 present at glutamatergic synapses, was moderately enhanced while gephyrin, present at GABAergic synapses, was massively enhanced following ictal-like and interictal-like EA (Fig. 6.D and E). Since PSD-95 was described as a lipid raft nucleating protein (Tulodziecka et al., 2016), and plasma membrane cholesterol depleting is known to reduce PSD-95 targeting to neuronal dendrites (Hering et al., 2003), these results are in conflict with previous observations reporting that enhanced excitatory synaptic transmission causes cholesterol depleting (Sodero et al., 2011), with evidence in this work that lipid rafts at synapses are destabilized by EA and with reports that synaptic AMPA receptor levels, known to be stabilized at synapses by palmitoylated PSD-95 (Han et al., 2015; Matt et al., 2019), are reduced following EA. However, this may reflect a specific response of the postsynaptic active zone to EA, since maintaining postsynaptic lipid rafts may be crucial to prevent epileptogenesis and abnormal spine growth by keeping together the molecular machinery allowing for AMPA receptor depalmitoylation and internalization (Fig. 7) (Han et al., 2015; Itoh et al., 2018).

GABAergic transmission and interneuron networks are also important players in seizure initiation and in epileptogenesis (Magloire et al., 2019; Avoli et al., 2022). At postsynaptic sites, GABA_A_ receptor recruitment to inhibitory GABAergic synapses requires the scaffold protein gephyrin and the action of the guanine nucleotide exchange factor collybistin through a PI3P dependent-process (Papadopoulos et al., 2017). Collybistin mutants with reduced lipid binding affinity disrupt GABAergic synapses by preventing gephyrin synaptic clustering and GABA_A_ receptor targeting and this was in turn reported to trigger epilepsy (Papadopoulos et al., 2015). Gephyrin postsynaptic distribution is organized in a lattice-like fashion and constitutes a dynamic puzzle of gephyrin nano-domains with different size and density (Pizzarelli et al., 2020). In addition, gephyrin is also regulated by palmitoylation, facilitating its membrane insertion and macromolecular clustering that also facilitates GABAergic transmission (Dejanovic et al., 2014; Matt et al., 2019). As such, the enhanced synaptic levels of gephyrin found in this work may reflect its synaptic recruitment to reinforce GABAergic transmission, through GABA_A_ recruitment, as an endogenous neuroprotective effort against hyperexcitability during ictal-like and interictal-like EA (Fig. 7) (Barberis, 2020). A similar response of GABAergic synapses was observed following anoxia (Lushnikova et al., 2011). Finally, our observations also contrast with the decreased levels of PSD-95 and gephyrin found in animal models and human MTLE (Sun et al., 2009; Fang et al., 2011; Bento-Oliveira, 2018), reinforcing the idea that acute and chronic synaptic responses to seizures are distinct.

In conclusion, LTP expression is impaired following EA, and this impairment is stronger following ictal-like than after interictal-like activity. Altered LTP expression is accompanied by marked modifications in several synaptic proteins involved in synaptic plasticity mechanisms such as AMPA receptor composition and phosphorylation and Kv4.2 levels. Furthermore, we observed altered synaptic membrane structure and domain regulating proteins like caveoilin-1, flotilin-1, PSD-95 and gephyrin. Altogether, these results suggest that early changes in synaptic plasticity are a promising target for antiepileptogenic therapies and strategies focusing on modulation of synaptic lipid raft domain dynamics may prove relevant to prevent epileptogenesis.

## Conflict of interests

The authors declare that the research was conducted in the absence of any commercial or financial relationships that could be construed as a potential conflict of interest.

## Ethical considerations

Animal housing and handing was performed in accordance with the Portuguese law (DL 113/2013) and European Community guidelines (86/609/EEC). The study was conducted according to the guidelines of the Declaration of Helsinki and approved by the Ethics Committee of the Faculty of Medicine, University of Lisbon.

## Author contributions

***JD Carvalho-Rosa:*** formal analysis and methodology; ***NC Rodrigues:*** formal analysis and methodology; ***A Silva-Cruz:*** formal analysis and methodology; ***SH Vaz:*** methodology; ***D Cunha-Reis:*** formal analysis and methodology, resources, supervision, funding acquisition, project administration, and writing – original draft, review, and editing.

## Funding

This work was supported national and international funding managed by Fundação para a Ciência e a Tecnologia (FCT, IP), Portugal. **Grants:** FCT UIDB/04046/2020 and UIDP/04046/2020 to BioISI, PTDC/SAU-NEU/103639/2008; and FCT/POCTI (PTDC/SAU-PUB/28311/2017) EPIRaft grant to DCR. **Fellowships:** SFRH/BPD/81358/2011 to DCR and **Researcher contract:** Norma Transitória - DL57/2016/CP1479/CT0044 to DCR.

## Acknowledgements

The authors wish to acknowledge the animal housing facilities of Instituto de Fisiologia, Faculdade de Medicina de Lisboa, and Laia Amat-Garcia and Cidália Gaspar for technical contribution.

## Data availability

The data that support the findings of this study are available from the corresponding author upon reasonable request. Some data may not be made available because of privacy or ethical restrictions.

## Abbreviations

0Mg^2+^: modified aCSF containing 0Mg2+ and 6mM elevated K^+^
aCSF: artificial cerebrospinal fluid
AEDs: antiepileptic drugs
Bic: bicuculine
EA: epileptiform activity
MTLE: mesial temporal lobe epilepsy
TBS: theta-burst stimulation

## Notes

### Competing Interest Statement

The authors have declared no competing interest.

